# Cyr61 delivery promotes angiogenesis during bone fracture repair

**DOI:** 10.1101/2024.04.05.588239

**Authors:** Annemarie Lang, Emily A. Eastburn, Mousa Younesi, Madhura Nijsure, Carly Siciliano, Annapurna Pranatharthi Haran, Christopher J. Panebianco, Elizabeth Seidl, Rui Tang, Eben Alsberg, Nick J. Willett, Riccardo Gottardi, Dongeun Huh, Joel D. Boerckel

## Abstract

Compromised vascular supply and insufficient neovascularization impede bone repair, increasing risk of non-union. Cyr61, Cysteine-rich angiogenic inducer of 61kD (also known as CCN1), is a matricellular growth factor that is regulated by mechanical cues during fracture repair. Here, we map the distribution of endogenous Cyr61 during bone repair and evaluate the effects of recombinant Cyr61 delivery on vascularized bone regeneration. In vitro, Cyr61 treatment did not alter chondrogenesis or osteogenic gene expression, but significantly enhanced angiogenesis. In a mouse femoral fracture model, Cyr61 delivery did not alter cartilage or bone formation, but accelerated neovascularization during fracture repair. Early initiation of ambulatory mechanical loading disrupted Cyr61-induced neovascularization. Together, these data indicate that Cyr61 delivery can enhance angiogenesis during bone repair, particularly for fractures with stable fixation, and may have therapeutic potential for fractures with limited blood vessel supply.

## Introduction

Compromised vascular supply and insufficient neovascularization are primary clinical challenges to bone repair and regeneration. Osteoblasts require close vascular proximity for oxygen and nutrients (*1, 2*), and the osteoprogenitor cells that mediate bone repair mobilize via vascular invasion (*3*). In fractures, especially in bones that have low peripheral vascular supply, insufficient neovascularization impedes repair and elevates non-union risk (*4*). Insufficient vascularization is an even greater challenge for segmental bone defect regeneration, which suffers from both insufficient progenitor cell pools and an obliterated vascular bed (*5, 6*). Regenerative therapies that overcome these challenges for fracture healing and bone defect regeneration could be transformative.

Mechanical stimuli determine the mode of bone healing. While low interfragmentary strains promote intramembranous (direct) ossification, high strains induce endochondral ossification (through a cartilage callus) (*7-9*). Previously, we found that mechanical loading can either promote or enhance bone healing, depending on interfragmentary strain magnitude and timing. Further, we found that these mechanical cues directly regulate neovascularization during bone regeneration (*10-12*). Mechanistically, we identified the transcriptional regulator Yes-associated protein (YAP) and transcriptional co-activator with PDZ-binding motif (TAZ) as key mechano-transducers during fracture repair (*13*), bone development (*14, 15*) and angiogenesis (*12, 16*). However, YAP and TAZ are oncogenes (*17*), suggesting that targeted activation of YAP/TAZ themselves would not be a feasible therapeutic for bone repair. However, multiple studies from our lab (*11-13, 15, 16, 18*) and others (*19-22*) identify Cysteine-rich angiogenic inducer 61 (Cyr61, also known as CCN1), as a direct target of YAP/TAZ mechanosignaling that may direct regeneration.

Cyr61 is a matricellular growth factor that functions as an integrin ligand (*23-26*) and integrates into the matrix via its N-terminal heparin binding domain (*27, 28*). Cyr61, a product of a growth factor-inducible immediate early gene, is associated with the cell surface and the extracellular matrix (*27, 29, 30*), and has been reported to regulate both chondrogenesis and osteogenesis during skeletal development (*31-33*). Cyr61 has also been implicated in fracture repair. Specifically, Cyr61 expression is elevated by mechanical stimulation during fracture repair (*34, 35*), and single-nucleotide polymorphisms to Cyr61 in patients increases risk for fracture non-union (*36*). Only one study has delivered recombinant Cyr61 during bone repair, investigating distraction osteogenesis in a rabbit model (*37*). In this study, a Cyr61-coated collagen sponge was wrapped around the osteotomy site at the time of surgery, prior to distraction (*37*). Cyr61 delivery increased bone strength, but did not significantly alter bone volume, suggesting Cyr61 may not simply promote osteogenesis during bone repair (*37*). The cellular targets of Cyr61 delivery and impacts of Cyr61 presentation on mechanoregulation of bone repair are unknown.

We hypothesized that the mechano-activated YAP/TAZ target gene Cyr61 can promote bone regeneration by mediating angiogenic/osteogenic crosstalk. Here, we map the distribution of endogenous Cyr61 during bone repair and evaluate the effects of recombinant Cyr61 delivery, under varied ambulatory loading conditions, effected by stiff or compliant fixation. We found that endogenous Cyr61 associates with the vascularized extracellular matrix and cellular YAP abundance in endochondral tissues after fracture. Treatment with Cyr61 did not promote chondrogenesis or osteogenesis in vitro or in the fracture callus in vivo. In contrast, Cyr61 treatment induced endothelial tube formation and vessel maturation in vitro and promoted neovascularization in the fracture callus. However, early ambulatory mechanical loading abrogated the angiogenic effects of Cyr61 delivery. Together, we found that Cyr61 delivery promoted angiogenesis during fracture repair, but not under mechanical conditions that were mechanically unfavorable for functional vascularization.

## Results

### Cyr61 associates with vascularized extracellular matrix after fracture

First, to understand how exogenous Cyr61 delivery might influence fracture repair, we examined the spatial patterns of endogenous Cyr61 abundance, its upstream regulator, YAP, and their spatial relationship to the vasculature and osteo-/chondro-progenitor cells at 14 days post-fracture (dpf) in a mouse femoral fracture model (**Fig. 1A, B**). We stabilized fractures with either stiff or compliant fixators to alter ambulatory load transfer and interfragmentary motion (*38*).

**Figure 1.**
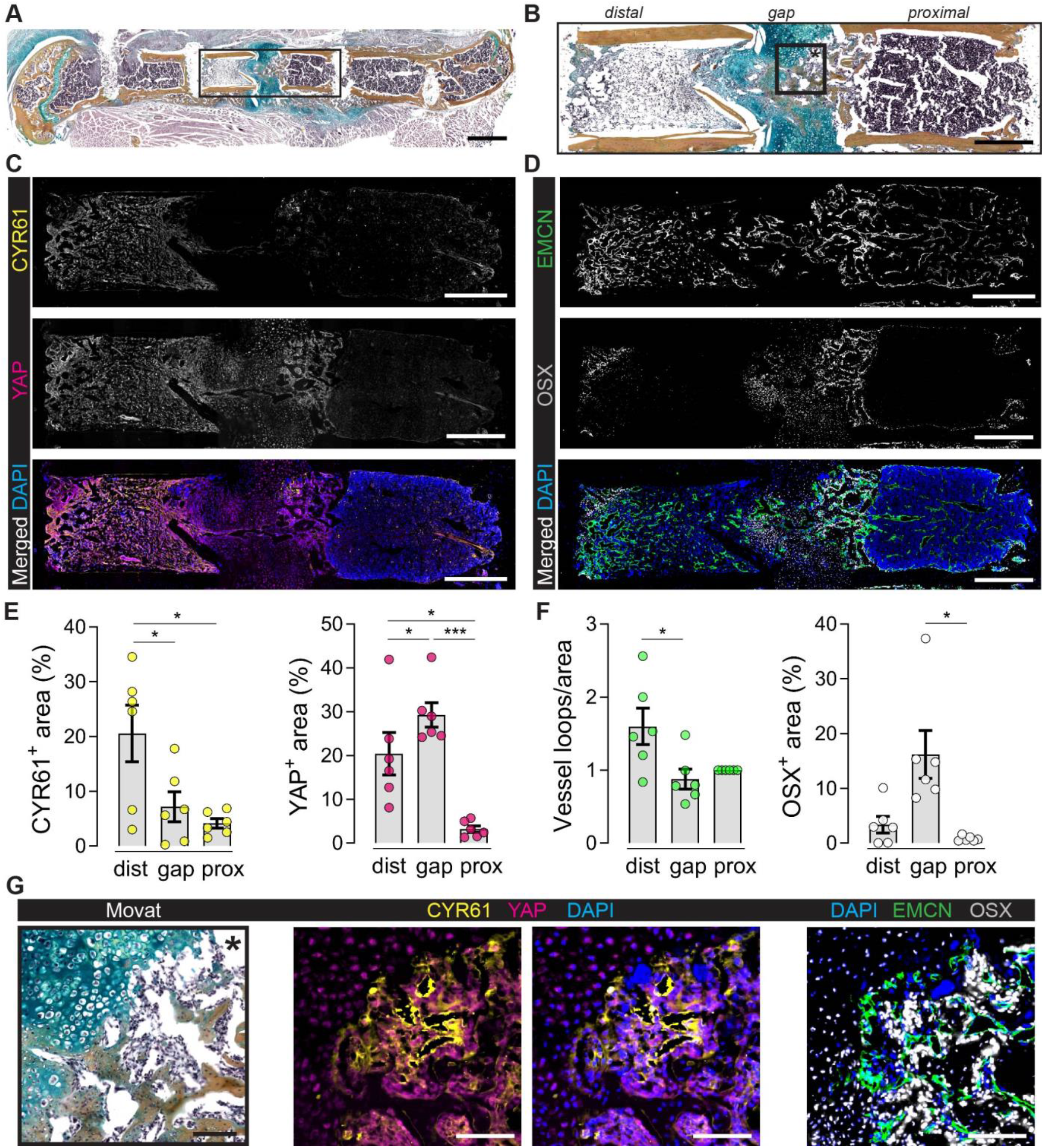
Spatial expression of YAP, CCN1/CYR61, EMCN and OSX in the fractured bone at 14 dpf. (**A**) Exemplary Movat’s Pentachrome staining 14 dpf; black box indicates ROI as magnified in B, C and D. (**B**) Magnification of fracture gap and adjacent bone marrow areas (ROI from A; black box indicates ROI for G). (**C, D**) Overview stainings. (**E, F**) Quantifications. One-way ANOVA with matched pairs was used to determine the statistical significance; p-values are indicated with *\*p < 0*.*05; ***p < 0*.*001*. (**G**) Magnifications from endochondral part of fracture gap. Scale bars indicate 1 mm (A) and 500 μm (B-D) and 100 μm (G).

We stained fractures for Cyr61, YAP, endomucin (EMCN, to mark endothelial cells) and osterix (OSX, to mark osteoprogenitor-lineage cells and hypertrophic chondrocytes) (**Fig. 1C, D**). Differences between stiff and compliant fixation on Cyr61 abundance and spatial distribution were not statistically significant (**Supplementary Fig. S1**). Therefore, for further image analysis, we combined these groups and defined three different regions of interest (ROI): the fracture gap (between the bone ends) and the distal and proximal bone marrow, spanning the endosteal region between the fracture gap and the next pin (**Fig. 1A**).

Cyr61 abundance was most prominent in the fracture-adjacent bone marrow (**Fig. 1C, E**), but was not highly expressed in the fracture gap (**Fig. 1G**) consistent with prior reports (*34*). Cyr61 staining was particularly evident in close proximity to EMCN-positive endothelial cells (**Fig. 1C-G**). Both OSX and YAP were abundant in the gap, consistent with our prior studies on the roles of YAP/TAZ signaling in OSX-expressing cells during endochondral bone regeneration and development (*13, 15*) (cf. **Fig. 1F**, Fig **1E**).

### Cyr61 enhances endothelial tube formation in vitro, but not hMSC chondrogenesis or osteogenesis

Next, to determine the direct effects of Cyr61 on chondrogenesis, osteogenesis and neovascularization, we evaluated human bone marrow stromal cells (hMSCs) differentiation and 3D angiogenesis in vitro (**Fig. 2A, E**). First, we evaluated the effects of exogenous Cyr61 on cartilage matrix deposition in a transforming growth factor (TGF)-β1-induced pellet chondrogenesis assay and osteogenic gene induction in an osteogenic medium differentiation assay (**Fig. 2A**). In the pellet assay, Cyr61 treatment modestly increased pellet size in a dose-dependent manner, in the absence of TGF-β1, but did not induce chondrocyte formation or glycosaminoglycan (GAG) deposition (**Fig. 2B**). Treatment with TGF-β1 markedly induced chondrogenesis and GAG deposition, but Cyr61 co-treatment had no effect on chondrogenesis, GAG production, or pellet size (**Fig. 2B, C**). In the osteogenesis assay, osteogenic medium significantly induced mRNA expression of osteogenic marker genes *RUNX2, SP7*, and *ALP*, as well as expression of *CYR61* (**Fig. 2D**). However, while exogenous Cyr61 treatment modestly, but significantly, increased *RUNX2* expression in osteogenic conditions, Cyr61 treatment did not alter *SP7* or *ALP* expression, or expression of Cyr61 itself (**Fig. 2D**).

**Figure 2.**
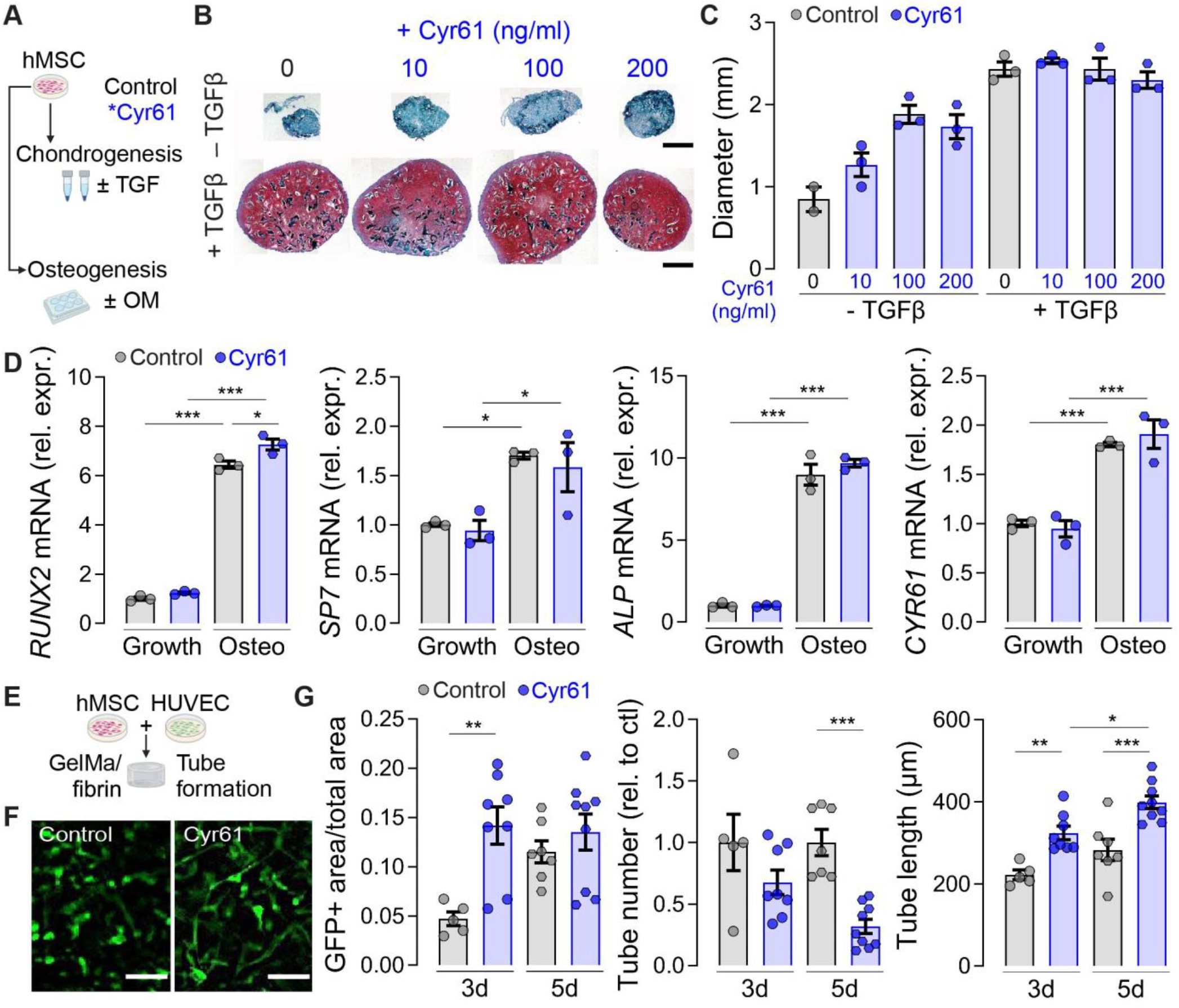
Cyr61 accelerates in vitro angiogenesis, but not chondrogenesis or osteogenesis of hMSCs. (**A**) Experimental setup. (**B**) Representative images of Safranin-O-staining and (**C**) quantitative measurement of chondrogenic pellet diameter at 2 weeks. (**D**) Relative mRNA expression of *RUNX2, SP7, COL1A1* and *CYR61* after 3 weeks of osteogenic differentiation normalized to housekeeping gene and control. (**E**) 3D in vitro angiogenesis assay combing HUVECs and hMSCs. (**F**) Exemplary images of tube formation at 3 days. (**G**) Quantification of relative GFP+ cell area, relative tube number and tube length. Mean ± SEM and individual data points. One-way ANOVA was used to determine the statistical significance; p-values are indicated with *\*p < 0*.*05; **p < 0*.*01; ***p < 0*.*001*. Scale bars indicate 500 μm (B) and 200 μm (F).

Next, we evaluated the effect of exogenous Cyr61 treatment on endothelial tube formation in a 3D in vitro angiogenesis assay featuring co-culture of GFP-labeled human umbilical vein endothelial cells (HUVECs) and hMSCs (**Fig. 2E**). Cyr61 treatment significantly increased GFP+ cell area density at 3 days (**Fig. 2G**). Cyr61-increased tubular network formation coincided with lower relative tube numbers at 3 and 5 days (**Fig. 2G**) and with increased tube length (**Fig. 2G**).

Together, these data suggest that Cyr61 has only modest effects on chondrogenesis and osteogenesis, under these in vitro conditions, but that Cyr61 stimulated tubular network formation.

### Local Cyr61 delivery had no effect on fracture callus bone or cartilage formation at 14 dpf

Next, to determine the effect of Cyr61 delivery on bone regeneration under varied ambulatory loading, we used a Gelatin methacrylate GelMA/fibrin scaffold to deliver Cyr61 to mouse femoral osteotomies, stabilized by either stiff or compliant fixation (**Fig. 3A**). The GelMA/fibrin scaffolds exhibited robust and persistent binding of Cyr61 in vitro (**Supplementary Fig. S2**). Neither fixation stiffness nor Cyr61 delivery significantly altered callus volume or bone volume at 14 dpf (**Fig. 3B, C**). Movat’s Pentachrome staining indicated more pronounced formation of fibrous tissue with Cy61 treatment in the rigid fixation group when compared to the non-treated group (**Fig. 3D**) with no significant differences in the relative mineralized bone or cartilage area (**Fig. 3E**).

**Figure 3.**
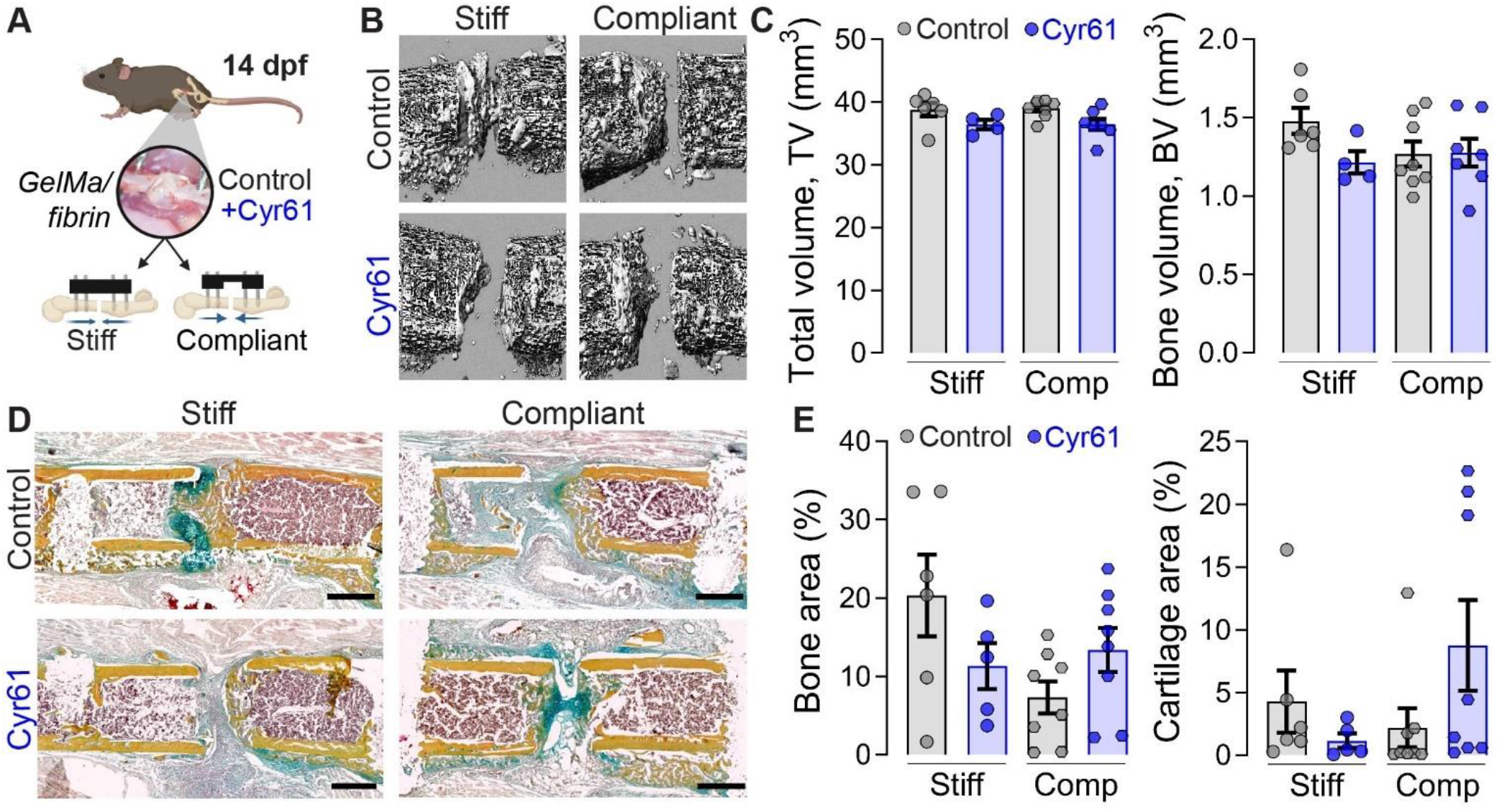
Cyr61 treatment had no effect on fracture callus bone or cartilage formation at 14 dpf. (**A**) Experimental setup. (**B**) 3D Representative reconstruction images of microCT analysis. (**C**) MicroCT - quantification of bone volume (BV) and bone volume fraction (bone volume/BV; total callus volume/TV). (**D**) Exemplary images of Movat’s Pentachrome staining – yellow = mineralized bone; green = cartilage; magenta = bone marrow; red = muscle tissue. (**E**) Quantification of mineralized bone and cartilage area in gap. Mean ± SEM and individual data points. One-way ANOVA was used to determine the statistical significance. Scale bars indicate 500 μm (D).

Taken together, we did not observe a significant effect of local Cyr61 delivery on bone or cartilage formation at 14 dpf.

### Local Cyr61 delivery strongly promoted vascular formation in callus area 14 dpf

Next, we asked whether Cyr61 delivery would promote neovascularization during fracture repair, depending on the mechanical environment. We analyzed immunofluorescence stainings using EMCN to mark blood vessels, OSX to mark osteoblast-lineage cells, F4/80 to mark macrophages and SOX9 to mark chondrocytes. Cyr61 significantly increased vessel formation under rigid fixation compared to vehicle control, but this effect was abrogated under compliant fixation (**Fig. 4A, B**). Cyr61 delivery did not significantly alter the amount or distribution of OSX+ cells (**Fig. 4A, B**). Macrophage invasion was qualitatively elevated by compliant fixation, and this was suppressed by Cyr61 delivery, though these differences were not statistically significant (**Fig. 4C, D**). No significant differences were observed in SOX9 staining (**Fig. 4C, D**).

**Figure 4.**
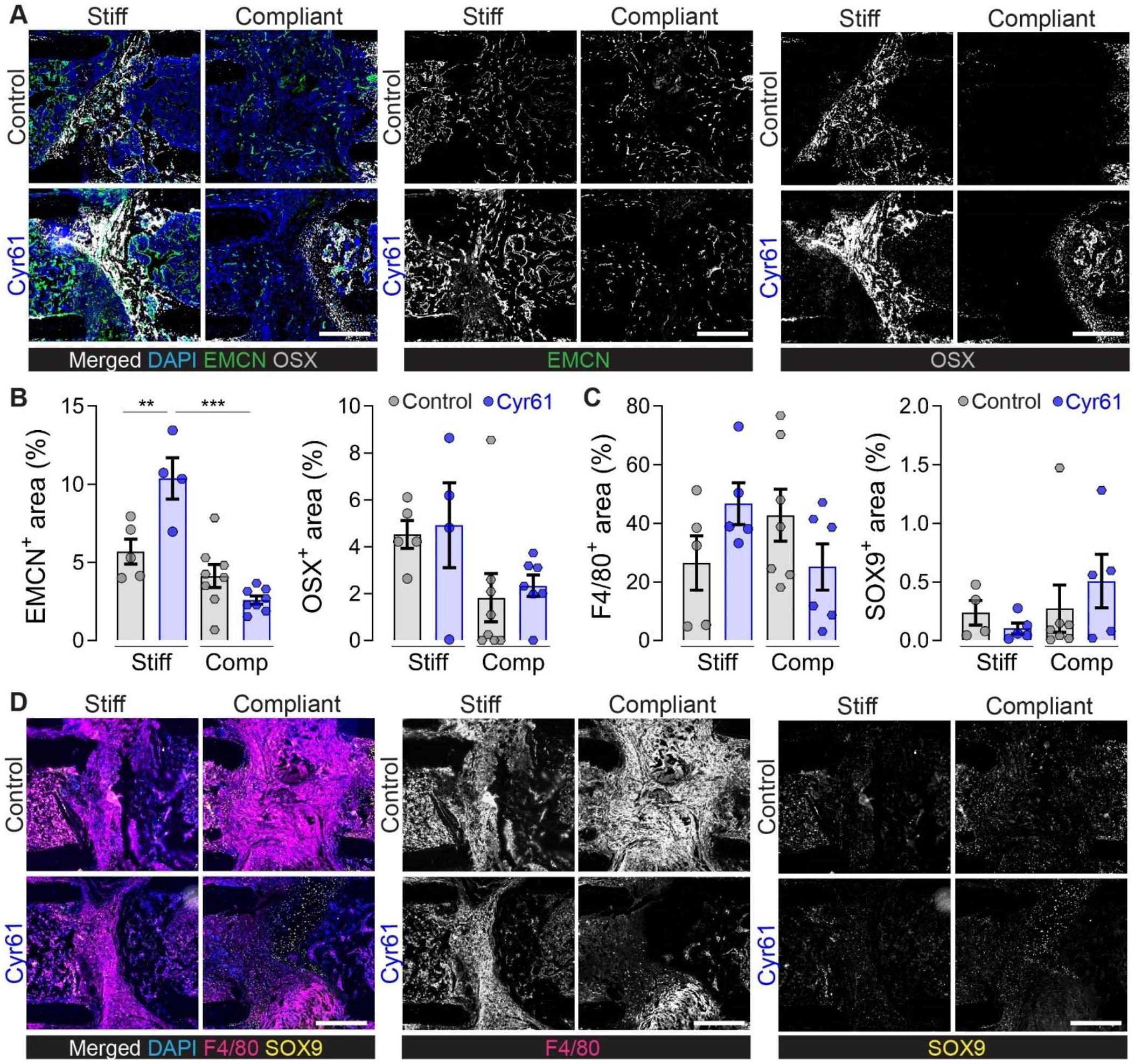
Treatment with endogenous Cyr61 promotes revascularization in the fracture gap 14 dpf. (**A**) Representative images of EMCN and OSX staining and (**B**) quantifications. (**C**) Quantifications and (**D**) representative images of F4/80 and SOX9 staining. Mean ± SEM and individual data points. One-way ANOVA was used to determine the statistical significance; p-values are indicated with *\*p < 0*.*05; ***p < 0*.*001*. Scale bars indicate 200 μm (A, D).

Together, these data suggest that Cyr61 delivery can promote angiogenesis during fracture repair, but this effect can be abrogated by initiation of ambulatory mechanical loading immediately after fracture.

### Cyr61 promotes vascular maturation with or without loading in a microphysiological vascularized bone-on-a-chip system

Previously, we found that mechanical loading regulates angiogenesis during bone regeneration, depending on the load magnitude and timing (*11, 39*). Specifically, we found that early loading disrupted neovascularization while delayed loading enhanced angiogenesis, both in vivo and in vitro. Since we observed here that early mechanical loading abrogated the angiogenic effect of Cyr61 treatment, we next sought to interrogate the interactions between the pro-angiogenic capacities of Cy61 and the timing of mechanical load initiation. To this end, we used a vascularized bone-on-a-chip system that combines hMSCs, human fibroblasts and endothelial progenitors embedded in a fibrin hydrogel on a microphysiological platform that allows for continuous perfusion with cell culture medium and dynamic mechanical loading. Compressive loading was applied at 10% compression for 1h per day, applied under two different loading scenarios, compared to a non-loaded static control: early loading (initiated at day 0, for 7 days) and delayed loading (initiated at day 4, followed by three days of loading) (**Fig. 5A**). Cyr61, or PBS control, were added to the cell culture medium from day 0. Samples were fixed and images were taken at 7 days (**Fig. 5B**).

**Figure 5.**
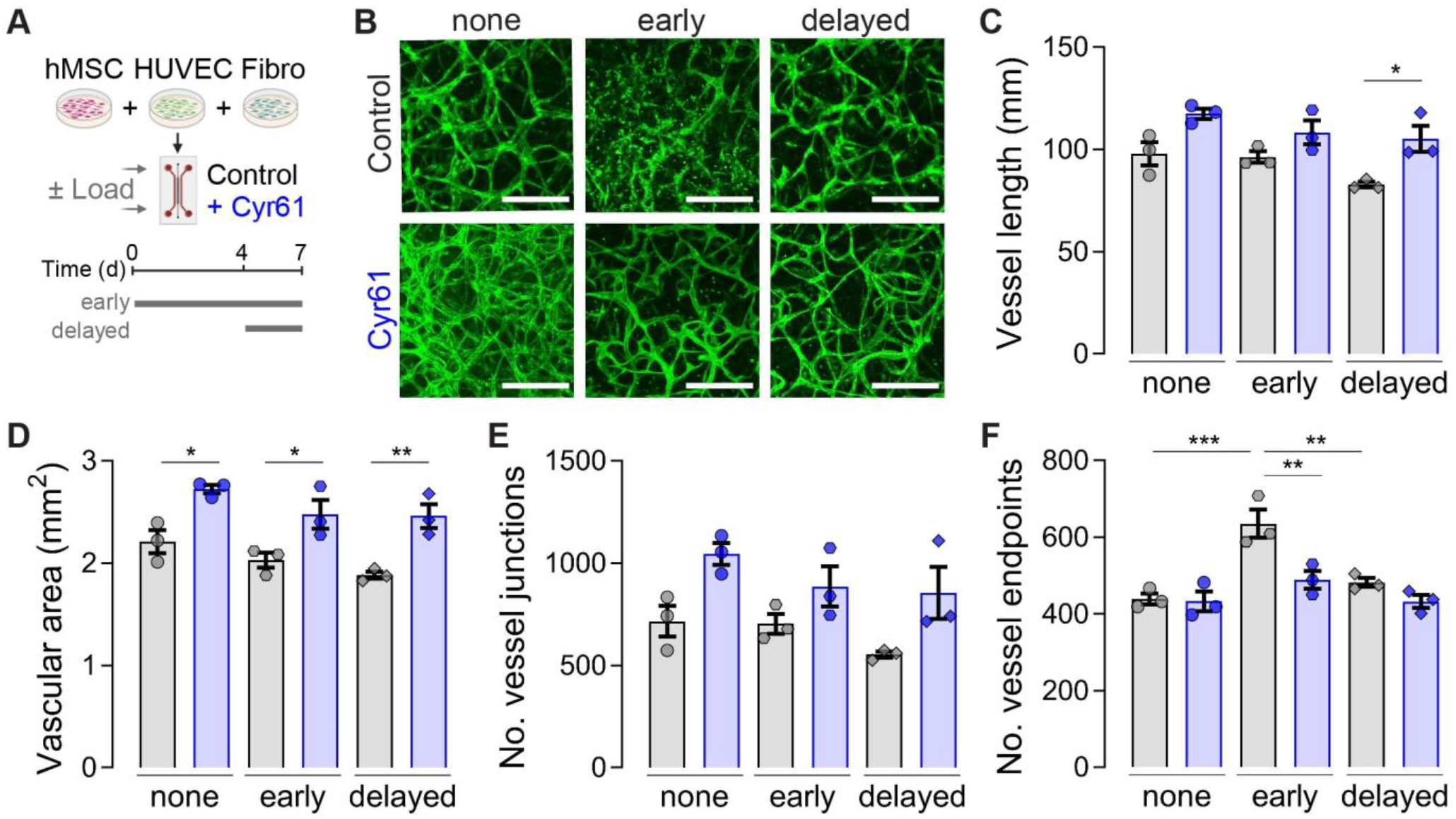
Cyr61 promotes vascular maturation with or without loading in vitro. (**A**) Experimental setup. (**B**) Representative images. (**C-F**) Quantifications of (**C**) vessel length, (**D**) vascular area, (**E**) number of vessel junctions and (**F**) number of vessel endpoints as measure for vascular connectivity. Absolute values are given for the whole chip area. Mean ± SEM and individual data points. One-way ANOVA was used to determine the statistical significance; p-values are indicated with *\*p < 0*.*05; **p < 0*.*01; ***p < 0*.*001*. Scale bars indicate 500 μm.

Cyr61 treatment significantly increased vessel length, only under delayed loading conditions, and increased vascular network area regardless of mechanical loading (**Fig. 5B-D**). Cyr61 treatment did not significantly alter vessel junction number (**Fig. 5E**). To assess network connectivity, we quantified the number of vessel endpoints. Early loading increased the vessel endpoint number, indicative of impaired network connectivity. This was rescued by Cyr61 treatment, indicating that early loading disrupted network formation, which was prevented by Cyr61 stimulation (**Fig. 5F**).

## Discussion

Here, we mapped the distribution of endogenous Cyr61 during bone repair and evaluated the effects of recombinant Cyr61 delivery, under varied ambulatory loading conditions, effected by stiff or compliant fixation. We found that endogenous Cyr61 associates with the vascularized extracellular matrix and cellular YAP abundance in endochondral tissues after fracture. Cyr61 treatment did not induce chondrogenesis or osteogenesis either in the fracture callus or in isolated cell culture in vitro. In contrast, Cyr61 treatment enhanced endothelial tube formation and maturation in vitro and promoted neovascularization in bone fracture; however, the angiogenic effects of Cyr61 treatment were abrogated by early mechanical loading.

### Effects of mechanical loading on angiogenesis during bone repair

Mechanical conditions at the fracture site determine the course and outcome of fracture repair, and the timing of mechanical load initiation is critical. Previously, we demonstrated that early ambulatory loading, which causes high interfragmentary strains, disrupts new blood vessel formation in the defect, while delayed loading profoundly enhances neovascularization (*11, 39*). Here, we found that early mechanical loading disrupts neovascularization, both during fracture repair in vivo and in vascularized tissue-on-a-chip experiments in vitro. These data are also consistent with our prior studies showing that dynamic matrix strain can directly alter 3D vascular structure formation by stromal vascular fragments cultured in vitro. In this prior study, we found that early application of dynamic matrix compression at high strain (30%) inhibited vessel formation, but delayed loading significantly increased both vessel length and branching (*12*). Further, in vitro mechanical strain induced robust Cyr61 expression in the stromal vascular composites and was abrogated by blockade of YAP/TAZ signaling (*12*). Together, these data support a profound mechanosensitivity of neovascular networks during fracture repair and point to Cyr61 as a potential mechano-responsive regulator of angiogenesis.

### Roles of YAP/TAZ mechanotransduction in skeletal development and repair

Previously, we and others have established the mechanoresponsive transcriptional regulators, YAP and TAZ, as critical mediators of mechanobiology (*19*) of both neovascularization (*40-42*) and endochondral ossification (*43*) during bone development (*14, 15, 18*) and fracture repair (*11, 13*). For example, conditional deletion of YAP and TAZ from Osx-expressing osteoblast-lineage cells impaired the co-mobilization of osteoprogenitors and blood vessels into the limb primary ossification center, disrupting vessel morphogenesis and vessel-mediated cartilage anlage remodeling (*14*). Further, using a fracture healing model (intramedullary pin fixation) that allows large interfragmentary strains and heals via endochondral ossification, we found that Osx-conditional YAP/TAZ deletion impaired vascularized fracture healing by regulating periosteal progenitor cell proliferation, osteoblastic differentiation, and osteogenic-angiogenic coupling, coincident with reduced Cyr61 expression (*13*). Together, these and other data identify YAP/TAZ mechanoactivation as a robust transcriptional mechanism for mechanoregulation of vascularized bone formation. However, due to their oncogenicity, YAP and TAZ themselves are not ideal targets for therapeutic activation (*44, 45*). Thus, targeting genes downstream of YAP/TAZ mechanosignaling may lead to safer and more efficient therapies for vascularized fracture healing. Here, we evaluated the effects of biomaterial-mediated delivery of recombinant Cyr61 on vascularized bone fracture repair.

### Cyr61 induces angiogenesis during bone repair

Our findings support prior studies showing that Cyr61 is upregulated during bone fracture repair and is induced by mechanical loading (*34, 35*). We found both YAP and Cyr61 to be abundant in the vascularized bone marrow, matrix adjacent to the fracture gap. This is consistent with the prior report from Hadjiargyrou and colleagues on Cyr61 expression during fracture repair, showing pronounced Cyr61 expression in vascularized fibrous tissues and periosteum (*34*). Likewise, Lineau et al. proposed a relationship between Cyr61 and angiogenesis during the early phase of fracture healing using an ovine model, showing that Cyr61 expression and vessel formation were impaired by early ambulatory mechanical loading (*35*).

Cyr61 promotes angiogenesis by stimulating endothelial cell migration and proliferation (*27, 46*). Cyr61 interacts with integrin receptors on the surface of endothelial cells, fibroblasts, and smooth muscle cells, triggering intracellular signaling pathways that promote cell motility and proliferation (*23-26*). We found that Cyr61 treatment increased tubular length and vessel maturation in vitro, consistent with a role for Cyr61 in promoting recruitment and assembly of endothelial cells into neovessel structures and facilitating their stabilization and maturation (*47*). Binding to cell surface receptors including as integrins and heparan sulfate proteoglycans, Cyr61 creates a chemotactic gradient that guides endothelial cells towards areas of tissue remodeling or injury where angiogenesis is required (*23-25, 27, 28*). Additionally, Cyr61 interacts with various growth factors and cytokines involved in angiogenesis, further enhancing its pro-angiogenic effects. For instance, Cyr61 potentiates the activity of vascular endothelial growth factor (VEGF)(*48, 49*), and can be activated by Plasminogen (*50*), to promote fracture repair (*51*). Consistent with these findings, we show that Cyr61 delivery robustly induces angiogenesis during fracture repair. These new data provide a mechanistic basis for the findings of Frey et al., who found that soluble Cyr61 delivery during distraction osteogenesis increased regenerated bone strength, without affecting callus formation or bone volume (*37*).

### Effects of Cyr61 on chondrogenesis and osteogenesis

Endogenous Cyr61 has been reported to mediate chondrogenesis during development due to its expression in both pre-chondrogenic mesenchyme and developing chondrocytes. (*31, 52*). Similarly, endogenous Cyr61 has been reported to mediate bone formation during development (*32, 33*). These studies show roles for Cyr61 in promoting chondrocyte proliferation and regulating key transcription factors involved in chondrogenesis and osteogenesis, such as Sox9 (*31*) and Runx2 (*53*). Cyr61 has also been reported to interact with various growth factors and signaling pathways known to regulate chondrogenesis and osteogenesis, including TGF-β, BMPs, and Wnt signaling, thereby orchestrating a complex network of molecular interactions critical for proper cartilage maturation (*31, 54, 55*). In this study, we did not observe significant effects of exogenous Cyr61 treatment on chondrogenesis or osteogenesis. Cyr61 treatment did not alter hMSC chondrogenic differentiation in vitro or on cartilage formation in the fracture callus in vivo. We observed a significant increase in *RUNX2* expression upon addition of Cyr61 during hMSC osteogenic differentiation in vitro but found with no effects on osteogenic commitment markers (*SP7, COL1A1, ALP*) or bone formation in the fracture callus at 14 dpf. Our data are consistent with the observations of Frey et al., who delivered soluble Cyr61 for distraction osteogenesis and likewise did not observe effects on bone volume, despite increases in regenerated bone strength (*37*).

Together, we found that Cyr61 delivery promoted angiogenesis during fracture repair, but this effect was abrogated by early mechanical loading. Thus, while potently angiogenic, both in vitro and in vivo, Cyr61 did not enhance vessel formation under mechanical conditions that were mechanically unfavorable for functional vascularization.

### Limitations

Cyr61 is a matricellular growth factor, but how its solubility vs. matrix tethering impacts signaling activity in vivo remains an active area of interest (*23, 27, 50, 51*). Here, we used a biomaterial system for Cyr61 delivery that did not completely remodel during the course of the experiment, as indicated by histological imaging. Further, our in vitro release data suggest that, consistent with its matrix-binding role, the amount of exogenous Cyr61 released from the fibrin matrix is modest. Thus, cellular interaction, perhaps through integrin adhesions, or cell-mediated matrix degradation may be necessary for the full beneficial effects of Cyr61 treatment. The degree of matrix entrapment or release may also influence the identity and response of effector cells. Based on our data, we postulate that Cyr61 delivery represents a potential pro-angiogenic therapeutic for vascularized bone healing under conditions of limited blood vessel supply, such as tibial fractures and large bone defects. However, our femoral fracture model is readily vascularized and heals spontaneously, so future studies in more challenging testbeds are warranted to evaluate the efficacy of Cyr61 delivery under stimulatory and vessel-supporting mechanical loading conditions.

## Materials and Methods

### Chondrocyte pellet generation and analysis

Human mesenchymal stromal cells (hMSCs) were isolated from bone marrow of healthy donors using a protocol approved by the University Hospitals of Cleveland Institutional Review Board (*11*). hMSCs were cultivated and expanded in growth media containing DMEM-LG (low glucose), 1% v/v penicillin-streptomycin (both Thermo Fisher Scientific, Waltham, MA), 10% v/v fetal bovine serum (Sigma Aldrich, St. Louis, MO) and 10 ng/ml fibroblast growth factor-2 (R&D Systems, Minneapolis, MN). For pellet cultures, hMSCs were detached, mixed with microspheres (0.75 mg/10^6^ cells) and transferred into conical V bottom plates (0.25 × 10^6^ cells/well), centrifuged for 5 min at 500 × *g* and incubated at 37 °C in a humidified incubator with 5% CO_2_ and 5% O_2_ (hypoxic conditions). Chondrogenic differentiation medium contained: DMEM-HG (high glucose), 1% v/v penicillin-streptomycin, 1% v/v ITS+ Premix Universal Culture Supplement (Corning, NY), 1 mM sodium pyruvate, 100 μM non-essential amino acids (both Thermo Fisher Scientific, Waltham, MA), 0.13 mM L-ascorbic acid-2-phosphate and 100 nM dexamethasone (both Sigma Aldrich, St. Louis, MO). Different concentrations of endogenous Cyr61 (0-200 ng/ml; Recombinant Human Cyr61/CCN1 Fc Chimera Protein, carrier-free; catalog no.: 4055-CR; R&D Systems, Minneapolis, MN) and TGFβ (0 or 10 ng/ml; rhTGF-beta1; R&D Systems, Minneapolis, MN) were added to the medium. Medium was changed every 3 days. Pellets were cultured for 14 days before being fixed with 4% paraformaldehyde (PFA; Electron Microscopy Sciences, Hatfield, PA) overnight at 4 °C, before switching to 70% ethanol and being paraffin embedded. Sections were stained with Safranin-O, images were taken with a Axio Observer (Carl Zeiss Microscopy, Wetzlar, Germany) and diameter measurement was performed using ImageJ.

### Osteogenic assay and RNA analysis

For osteogenic differentiation, hMSC were plated in a 6-well plate (3 × 10^5^ cells/well) and expanded until being confluent. Osteogenic differentiation was induced with growth medium supplemented with 10 mM β-glycerophosphate, 100 μM ascorbic acid and 100 nM dexamethasone (all Sigma Aldrich, St. Louis, MO). Endogenous Cyr61 (100 ng/ml) was added to the medium. Cells were incubated for 3 weeks at 37 °C in a humidified incubator with 5% CO_2_ and approx. 18% O_2_. Medium was changed every 3 days. RNA was isolated using the Qiagen RNeasy Kit (Qiagen, Hilden, Germany) following the manufacture’s instruction. RNA concentration was determined using Nanodrop (Thermo Fisher Scientific, Waltham, MA) and 0.5 μg was used for cDNA synthesis using the High-Capacity cDNA Reverse Transcription Kit (Applied Biosystems, Waltham, MA). qPCR was performed with the SYBR Green PCR Master mix (Applied Biosystems, Waltham, MA) and custom-designed qRT-PCR primers (IDT, Coralville, IA). We used the following primer sequences:

*RUNX2* – TGGCTGGTAGTGACCTGCGGA (reverse); ACAGAACCACAAGTGCGGTGCAA (forward)

*SP7* – TGGGCAGCTGGGGGTTCAGT (reverse); TGGCTAGGTGGTGGGCAGGG (forward)

*ALP* – GCAGTGAAGGGCTTCTTGTC (reverse); CCACGTCTTCACATTTGGTG (forward)

*CYR61* – GGTTGTATAGGATGCGAGGCT (reverse); GAGTGGGTCTGTGACGAGGAT

*GAPDH* – GGCTGGTGGTCCAGGGGTCT (reverse); GGGGCTGGCATTGCCCTCAA (forward)

Gene expression was normalized to *GAPDH* and calculated as fold change using the comparative CT method to the control (growth medium and no Cyr61).

### Generation of GelMA/fibrin hydrogel for in vitro angiogenesis assay and in vivo Cyr61 delivery

Gelatin methacrylate (GelMA; 5% w/w) and 5 mg/ml fibrinogen were mixed and dissolved in a 0.2% w/w lithium phenyl-2,4,6-trimethylbenzoylphosphinate (LAP) solution previously dissolved in sterile saline (all products were purchased from Cellink, Gothenburg, Sweden). To ensure proper dissolution, the mixture was incubated overnight at 37 °C on a shaking plate and frozen at -20 °C for storage. Aliquots were thawed in a 37 °C water bath prior to every new experiment. Crosslinking of the hydrogel was conducted in 2-steps - thrombin-induced fibrin formation and UV-initiated photocrosslinking. Thrombin was dissolved in 40 mM calcium chloride (CaCl_2_, Sigma Aldrich, St. Louis, MO) solution at a concertation 0.7 g/l which was added at a 1:80 dilution to the GelMA/fibrin solution. A UV flashlight was used for photocrosslinking for 2.5 minutes.

### In vitro 3D tube formation assay

GFP^+^ HUVECs were purchased and cultured in human endothelial growth medium (EGM-SF1; cells and medium from Angio-Proteomie, Boston, AM) with 5% v/v FBS (Sigma Aldrich, St. Louis, MO) in cell culture flasks (Corning, Corning, NY) coated with autoclaved 0.2% w/w gelatin (Sigma Aldrich, St. Louis, MO). hMSCs were expanded in RoosterNourish™-MSC-XF (both RoosterBio, Frederick, MD). HUVECs were starved with EGM with 0.1% v/v FBS 12h prior to the begin of the experiment. For 3D construct formation, HUVECs and hMSCs were detached from the culture flasks independently washed with PBS and mixed at a concentration of 4:1 in the liquid GelMA/fibrin hydrogel already containing thrombin (*14, 56*). The hydrogel/cell mixture was immediately transferred into a disc sized, silicone mold with a diameter of 6 mm and a thickness of 2 mm (capacity for 50 microL). UV light was applied subsequently for 2.5 min. The mold was carefully removed, and the construct was transferred into a 12-well culture plate filled with EGM, 0.1% v/v FBS with or without 100 ng/ml Cyr61 (Recombinant Human Cyr61/CCN1 Fc Chimera Protein, carrier-free; catalog no.: 4055-CR; R&D Systems, Minneapolis, MN). Medium change was performed at day 3 and images were taken at 3 and 5 days using a Keyence BZ-X800 Fluorescence Microscope (Keyence, Itasca, IL). All images were blinded for group and treatment. The relative tube number, mean tube length and relative GFP^+^ vascular area (normalized to the total area) were analyzed using Fiji ImageJ. The area to be analyzed was defined as the full area of the hydrogel which was visible in the images. To correct for differences in analyzed area (ROI) dimension between the hydrogels, the tube number was normalized to the analyzed area (ROI). We used two different MSC lines combined with two different HUVEC lines with 1-3 replicates per MSC/HUVEC combination and condition (biological replicates n= 4; technical replicates total n= 5-9).

### Local delivery system and Cyr61 release kinetic

A total of 1 μg Cyr61 (Recombinant Human Cyr61/CCN1 Fc Chimera Protein, carrier-free; catalog no.: 4055-CR; R&D Systems, Minneapolis, MN) was dissolved in GelMA/fibrin hydrogel containing thrombin and transferred into a cylinder-shaped mold with a liquid capacity of 8.6 μl and 1 mm diameter with 1 mm thickness. UV light was applied for 2.5 min and the solid hydrogels were removed from the mold with a fine forceps. As control, we generated the hydrogel following the same procedure without addition of Cyr61. For release kinetic measurement, hydrogels were produced as described before and cultivated for 14 days in 1 mL PBS at 37 °C. The PBS was completely collected and changed at 1h, 3h, 7h, 1d, 3d, 7d and 14d. The experiment was performed in a triplet and samples were frozen at -20 °C until further use. Cyr61 concentration was determined using a human Cyr61 Quantikine ELISA (R&D Systems, Minneapolis, MN) following the manufacturer’s instructions.

### Animals and surgical procedure

A total of 32 C57BL/6J female mice (Charles River Laboratories, Wilmington, MA) aged 12-16 weeks underwent surgery. All procedures were conducted in accordance with IACUC regulations (University of Pennsylvania; protocol no: 806482). Veterinary care and animal husbandry was provided by University Laboratory Animal Resources (ULAR) at the University of Pennsylvania in accordance with contemporary best practice.

Mice were housed in a semi-barrier facility in cages (Ancare Corp., Bellmore, NY). Housing conditions encompassed a 12/12–h light/dark cycle (light from 7:00 a.m. to 7:00 p.m.), room temperature of 72 ± 2F and a humidity of 50 ± 10%. Food (Rodent Diet, LabDiet) and tap water were available *ad libitum*. Mice were randomly divided into pairs per cage. Cages contained wooden chips (Bed-o’Cobs 1/4, Laboratory Animal Bedding), Enviro-dri (Shepherd Specialty Papers, Milford, NJ), and a shredded paper towel as bedding and nesting material. Additional enrichment was provided such as a clear mouse transfer tube (Braintree Scientific, Braintree, MA), a mouse double swing (Datesand Group, Bredbury, United Kingdom) and a Shepherd Shack (Shepherd Specialty Papers, Milford, NJ) where the entrance area was enlarged to avoid injuries due to the external fixator (*57*). Transfer tube and double swing were removed after surgery to reduce the risk of injury. Animals were handled with the transfer tube.

Mice were anesthetized with isoflurane (∼2–3%; provided in 100% oxygen; Dechra Veterinary Products, Overland Park, KS) and moved onto a heating pad (37 °C; Kent Scientific, Torrington, CT). Anesthesia was maintained at ∼1.5–2% with an individual a nose cone. Eye ointment (Optixcare eye lube, Aventix, Ontario, Canada), physiological saline (0.9% sodium chloride; 0.5 ml, s.c.; BD, Franklin Lakes, NJ), clindamycin (45 mg/kg, s.c.; Sagent Pharmaceuticals, Schaumburg, IL) and Buprenorphine SR-Lab (1 mg/kg, s.c.; Wedgewood Pharmacy, Swedesboro, NJ) were applied. The left femur was shaved and disinfected with alcoholic iodine solution and 70% ethanol. A longitudinal skin incision was made between knee and hip. The musculus vastus lateralis and musculus biceps femoris were bluntly separated and the femur was exposed. Two different external fixators were used to mimic 2 distinct loading scenarios (stiff: 18.1 N/mm; compliant: 3.2 N/mm, both RISystem, Davos, Switzerland). The external bar of the fixator was positioned parallel to the femur and the pins were screwed into the bone after holes have been pre-drilled. An approximately 0.7 mm fracture/osteotomy gap was created between the second and third pin using a Gigli wire saw (0.66 mm; RISystem, Davos, Switzerland) and the gap was flushed with saline. The GelMa/fibrin scaffold with or without Cyr61 was placed between the bone ends. Muscle and skin were closed with two layers of sutures (muscle: coated Vicryl; skin: Prolene, both Ethicon, Raritan, NJ). The wound was covered with a triple antibiotic cream (B.N.P. Triple Antibiotic Ophthalmic Ointment, Neobacimyx-H, Schering Plough, Kenilworth, NJ). Mice were returned to their home cages placed under an infrared lamp and closely monitored until fully recovered. To ensure food and water uptake after surgery, Diet Gel (ClearH_2_O, Westbrook, ME) was provided on the cage floor.

Mice were monitored closely and scored during the first 4 days, at day 7 and day 10 before being euthanized at day 14. The scoring sheet was based on a composite score consisting of the mouse grimace score (eyes and ears only), a clinical scire and a model specific score including limping and dragging score following previous established systems (*58, 59*). Humane endpoints were defined before the experiment and included: wound suture completely discreet, no coat care/feces soiling, sunken/glued eyes, hunched back, periprosthetic fracture, gross malposition >20° axial deviation of the fracture ends, no food and water intake and weight loss >25%, bloody feces, significantly increased breathing/wheezing, diarrhea (if debilitating or persistent), seizures/staggering/apathy, paresis of more than two limbs and abscesses. No humane endpoint was reached during the study.

Mice were euthanized 14 days after surgery using CO_2_ and cervical dislocation. The fractured femora were collected and fixed in 4% PFA at 4 °C for 7h and transferred to PBS until ex vivo microCT was completed.

All analyses were performed after samples were blinded for groups (fixation, treatment). De-blinding was performed once all samples were analyzed to avoid any bias.

### Ex vivo microCT

Ex vivo microCT was performed using a μCT 45 desktop scanner (Scanco Medical AG, Brüttisellen, Switzerland) after removal of the external fixator and fixation of the bones in plastic pipettes to avoid destruction of the callus tissue. The area in between the inner two pins was scanned with an isotropic voxel size of 10.4 μm (55 kVp, 72 μA, AL 0.5 mm, 1x400 ms) and the scan axis was aligned along the diaphyseal axis of the femora. 3D reconstruction and analyses were performed using the provided software package (global threshold of 240 mg HA/cm^3^) analyzing a fixed VOI (200 slices total) starting from the middle of the fracture gap. The fixed VOI was transferred to every sample to allow for a standardized total volume across samples. The original cortical bone was excluded to only analyze newly formed bone.

### Histology and immunofluorescence

Following microCT, femora were then transferred into 10% w/v ethylenediaminetetraacetic acid (EDTA) pH 7.4 for 3 days at 4 °C and transferred to 30% w/v sucrose solution for 2 days before being cryo-embedded using Tissue-Tek OCT (Sakura Finetek USA, Torrance, CA). Consecutive sections of 7 μm were prepared (cyrotome, Leica, Wetzlar, Germany) using cryotape (Sectionlab, Japan). Sections were fixed onto glass slides and stored at -20 °C until staining. Movat’s pentachrome staining was performed using a ready to use kit (Morphisto, Offenbach am Main, Germany). The manufacturer’s protocol was adapted to cryo-sections based on previously used protocols (*58*). Imaging was performed on a AxioScan (Carl Zeiss Microscopy, Wetzlar, Germany) and quantitative analyses of the Movat’s pentachrome staining were evaluated using an ImageJ.

For immunofluorescence, sections were rehydrated in PBS. Blocking solution (10% v/v goat serum/PBS; Sigma Aldrich, St. Louis, MO) was added for 30 min and antibodies were diluted in PBS/0.1% v/v Tween20/5% v/v goat serum (Sigma Aldrich, St. Louis, MO) or PBS/3% v/v Triton/5% v/v goat serum (YAP/Cyr61 only). The following primary antibodies and secondary antibodies were used (staining durations provided): CYR61 (R&D Systems, Minneapolis, MN; catalog number: MAB4864; 1:20; overnight at 4 °C), YAP (Cell Signaling, Danvers, MA; clone D8H1X; catalog number: 14074; 1:100; overnight at 4 °C), EMCN (Santa Cruz, Dallas, TX; clone V.5C7; catalog number: sc-65495; 1:100; 2h at RT), OSX (Abcam, Cambridge, United Kingdom; catalog number: ab209484; 1:100; 2h at RT), F4/80 (Novus Biological, Littleton, CO; Cl:A3-1, catalog number: NBP2-81030; 1:400; 2h at RT), Sox9 (Abcam, Cambridge, United Kingdom; catalog number: ab185230; 1:200; 2h at RT); all secondary antibodies were purchased from Thermo Fisher Scientific and used at an 1:500 dilution for 2h at RT if not stated otherwise: goat anti-rat A647 (A-21247), goat anti-rat A488 (A-11006), goat anti-rabbit A647 (A-27040), goat anti-rabbit A488 (Abcam; ab150077; 1:1,000). DAPI (NucBlue Fixed Cell ReadyProbes Reagent; Thermo Fisher Scientific, Waltham, MA) was added during the last washing step and sections were covered with Fluoromount-GT (Thermo Fisher Scientific, Waltham, MA). Images were taken with an AxioScan and image quantification was performed using the Fiji/ImageJ software. The area of interest was manually assigned with the built-in ROI-Manager and determined with the thresholding tool.

### Loading experiments with vascularized bone-on-a-chip system

To create the vascularized bone-on-a-chip system, 10 μl of fibrin gel containing fibrinogen at a final concentration of 10 mg/ml (F8630, Sigma Aldrich, St. Louis, MO) mixed with 1 U/ml thrombin (T7513, Sigma) was injected through the inlet access port into the middle lane of the culture chamber. The following cells were dispensed in the fibrin gel before injection into the microdevice: HUVECs (3.5 × 106 cells/ml), MSCs (2.5 × 10^6^ cells/ml) and human fibroblasts (2.5 × 10^6^). The microdevice was placed in a cell culture incubator to induce fibrin gelation at 37 °C for 10 minutes. Upon gelation, complete endothelial cell growth media was supplemented to the side channels of the culture chamber through media reservoirs. The side channels were seeded after 24h with endothelial cells (5 × 10^6^ cells/ml) to form endothelial lining on the channel walls and anastomosis of side channel with vasculature in hydrogel. Media (EGM-2; Lonza, Basel, Switzerland) in the reservoirs was changed every other day during the subsequent culture period. To investigate the effect of Cyr61 on vascular formation under mechanical stimuli, we examined 4 groups: without and with Cyr61 and/or no mechanical loading, mechanical loading from the beginning for 7 days and delayed loading starting at day 4. The microdevices were mechanically stimulated with a uniaxial compression load up to 10% strain with a frequency of 1 Hz (1 hour of mechanical stimuli each day). Samples were fixed at day 7 of culture and stained with CD31 human antibody (Abcam; ab134168; AF488-conjugated; 1:100) and imaged with an Inverted confocal microscope. Images were analyzed with AngioTool software (AngioTool64 0.6a) to quantify changes in vessel length, area, average vessel diameter and junction number. Analyses are performed for the whole chip area (fixed ROI: 7 mm^2^) and provided as absolute measures.

### Statistical analysis

GraphPad Prism V.9 was used for statistical analysis. Data was tested for Gaussian distribution according to D’Agostino-Pearson omnibus normality test and homoscedasticity. One-way ANOVA was used to determine the statistical significance. A p value <0.05 was considered statistically significant. Sample sizes are indicated in the graph displaying individual data points. Data are displayed with error bars showing mean ± SEM. All analyses were performed on distinct samples. Samples or data were only excluded in justified cases due to e.g. technical errors, or unrecognizable target structures.

## General

The authors would like to thank all members of the Boerckel lab for constructive discussions on the research presented in this study.

## Author contributions

A.L. and J.D.B. conceived and supervised the research. A.L., J.D.B., E.A.E., M.N., A.P.H., C.J.P., E.S. designed and performed animal experiments. A.L., E.A.E. and C.S. performed analysis. R.T. and E.A. provided materials and methods for in vitro chondrogenesis and osteogenesis assays. A.L. and R.G conceived and supervised in vitro angiogenesis assay. M.Y. and D.H. conceived and supervised experiments with the vascularized bone-on-a-chip system. A.L. and J. D.B. wrote the paper. All authors discussed data and revised the manuscript.

## Data and materials availability

All data are available in the main text or the supplementary materials.

## Supplementary Materials

**Figure S1.**
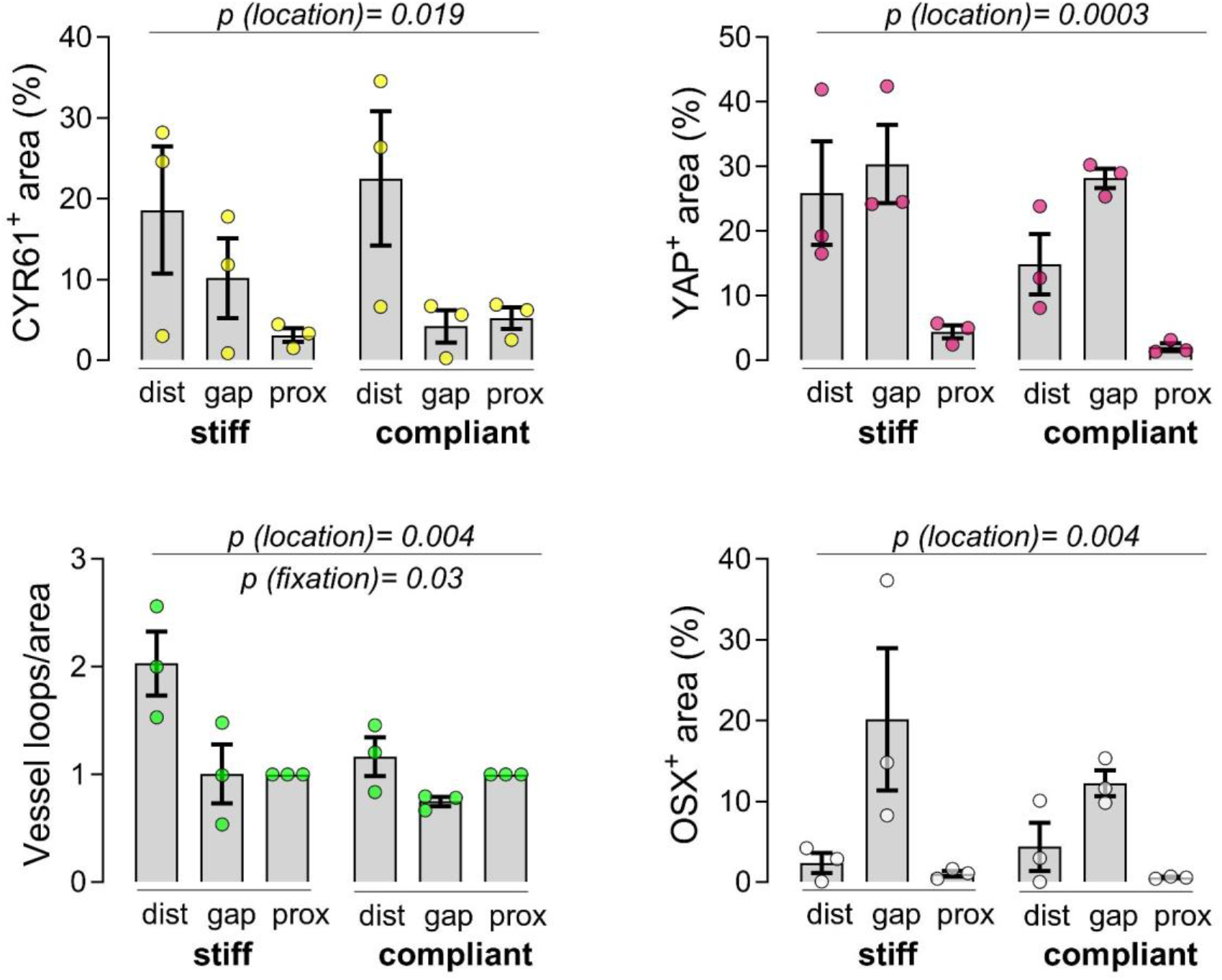
Spatial expression of YAP, CCN1/CYR61, EMCN and OSX stiff vs. compliant fixation. Separated quantifications. Ordinary two-way ANOVA was performed to determine main effects of location (distal, gap, proximal) and fixation (stiff vs. compliant). Statistical significances are provided in graphs.

**Figure S2.**
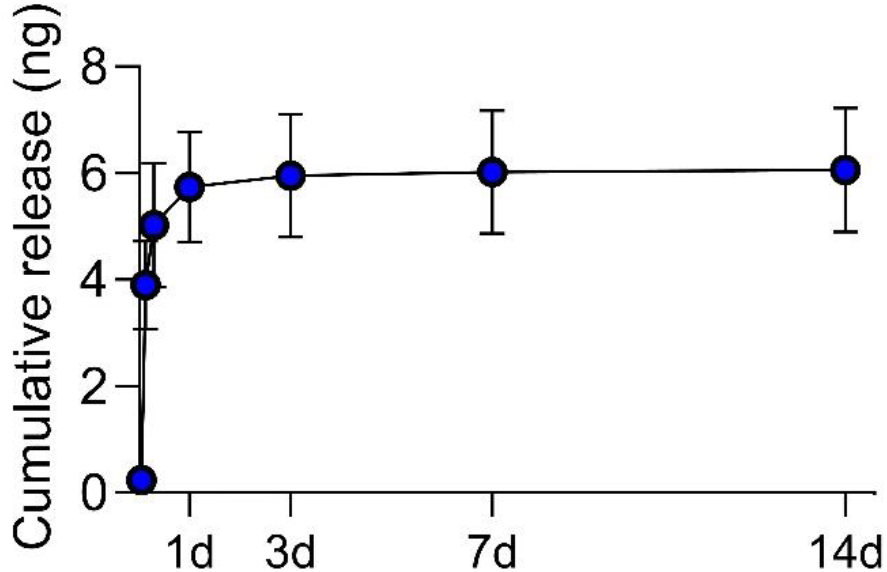
Cyr61 release kinetics from GelMA/fibrin scaffolds determined over 14d. 1 μg rhCyr61 was loaded into GelMA-fibrin scaffolds and release into 1 mL PBS was measured by ELISA over 14 days. Cumulative measured release averaged 6 ng (0.6% of total loaded protein).

## References

1. E. A. Phelps, A. J. García, Engineering more than a cell: vascularization strategies in tissue engineering. Curr Opin Biotechnol 21, 704–709 (2010).

2. N. van Gastel, S. Stegen, G. Eelen, S. Schoors, A. Carlier, V. W. Daniëls, N. Baryawno, D. Przybylski, M. Depypere, P. J. Stiers, D. Lambrechts, R. Van Looveren, S. Torrekens, A. Sharda, P. Agostinis, D. Lambrechts, F. Maes, J. V. Swinnen, L. Geris, H. Van Oosterwyck, B. Thienpont, P. Carmeliet, D. T. Scadden, G. Carmeliet, Lipid availability determines fate of skeletal progenitor cells via SOX9. Nature 579, 111–117 (2020).

3. C. Maes, T. Kobayashi, M. K. Selig, S. Torrekens, S. I. Roth, S. Mackem, G. Carmeliet, H. M. Kronenberg, Osteoblast precursors, but not mature osteoblasts, move into developing and fractured bones along with invading blood vessels. Dev Cell 19, 329–344 (2010).

4. J. Street, M. Bao, L. deGuzman, S. Bunting, F. V. Peale, Jr., N. Ferrara, H. Steinmetz, J. Hoeffel, J. L. Cleland, A. Daugherty, N. van Bruggen, H. P. Redmond, R. A. Carano, E. H. Filvaroff, Vascular endothelial growth factor stimulates bone repair by promoting angiogenesis and bone turnover. Proc Natl Acad Sci U S A 99, 9656–9661 (2002).

5. H. C. Fayaz, P. V. Giannoudis, M. S. Vrahas, R. M. Smith, C. Moran, H. C. Pape, C. Krettek, J. B. Jupiter, The role of stem cells in fracture healing and nonunion. International Orthopaedics 35, 1587–1597 (2011).

6. S. Stegen, N. van Gastel, G. Carmeliet, Bringing new life to damaged bone: the importance of angiogenesis in bone repair and regeneration. Bone 70, 19–27 (2015).

7. L. Claes, K. Eckert-Hübner, P. Augat, The effect of mechanical stability on local vascularization and tissue differentiation in callus healing. J Orthop Res 20, 1099–1105 (2002).

8. A. E. Goodship, J. Kenwright, The influence of induced micromovement upon the healing of experimental tibial fractures. J Bone Joint Surg Br 67, 650–655 (1985).

9. G. Matziolis, J. Tuischer, G. Kasper, M. Thompson, B. Bartmeyer, D. Krocker, C. Perka, G. Duda, Simulation of cell differentiation in fracture healing: mechanically loaded composite scaffolds in a novel bioreactor system. Tissue Eng 12, 201–208 (2006).

10. S. Herberg, A. M. McDermott, P. N. Dang, D. S. Alt, R. Tang, J. H. Dawahare, D. Varghai, J. Y. Shin, A. McMillan, A. D. Dikina, F. He, Y. B. Lee, Y. Cheng, K. Umemori, P. C. Wong, H. Park, J. D. Boerckel, E. Alsberg, Combinatorial morphogenetic and mechanical cues to mimic bone development for defect repair. Sci Adv 5, eaax2476 (2019).

11. A. M. McDermott, S. Herberg, D. E. Mason, J. M. Collins, H. B. Pearson, J. H. Dawahare, R. Tang, A. N. Patwa, M. W. Grinstaff, D. J. Kelly, E. Alsberg, J. D. Boerckel, Recapitulating bone development through engineered mesenchymal condensations and mechanical cues for tissue regeneration. Sci Transl Med 11, (2019).

12. M. A. Ruehle, E. A. Eastburn, S. A. LaBelle, L. Krishnan, J. A. Weiss, J. D. Boerckel, L. B. Wood, R. E. Guldberg, N. J. Willett, Extracellular matrix compression temporally regulates microvascular angiogenesis. Sci Adv 6, (2020).

13. C. D. Kegelman, M. P. Nijsure, Y. Moharrer, H. B. Pearson, J. H. Dawahare, K. M. Jordan, L. Qin, J. D. Boerckel, YAP and TAZ Promote Periosteal Osteoblast Precursor Expansion and Differentiation for Fracture Repair. J Bone Miner Res 36, 143–157 (2021).

14. J. M. Collins, A. Lang, C. Parisi, Y. Moharrer, M. P. Nijsure, J. H. Thomas Kim, S. Ahmed, G. L. Szeto, L. Qin, R. Gottardi, N. A. Dyment, N. C. Nowlan, J. D. Boerckel, YAP and TAZ couple osteoblast precursor mobilization to angiogenesis and mechanoregulation in murine bone development. Dev Cell, (2023).

15. C. D. Kegelman, D. E. Mason, J. H. Dawahare, D. J. Horan, G. D. Vigil, S. S. Howard, A. G. Robling, T. M. Bellido, J. D. Boerckel, Skeletal cell YAP and TAZ combinatorially promote bone development. Faseb j 32, 2706–2721 (2018).

16. D. E. Mason, J. M. Collins, J. H. Dawahare, T. D. Nguyen, Y. Lin, S. L. Voytik-Harbin, P. Zorlutuna, M. C. Yoder, J. D. Boerckel, YAP and TAZ limit cytoskeletal and focal adhesion maturation to enable persistent cell motility. J Cell Biol 218, 1369–1389 (2019).

17. J. M. Franklin, Z. Wu, K. L. Guan, Insights into recent findings and clinical application of YAP and TAZ in cancer. Nat Rev Cancer 23, 512–525 (2023).

18. C. D. Kegelman, J. C. Coulombe, K. M. Jordan, D. J. Horan, L. Qin, A. G. Robling, V. L. Ferguson, T. M. Bellido, J. D. Boerckel, YAP and TAZ Mediate Osteocyte Perilacunar/Canalicular Remodeling. J Bone Miner Res 35, 196–210 (2020).

19. S. Dupont, L. Morsut, M. Aragona, E. Enzo, S. Giulitti, M. Cordenonsi, F. Zanconato, J. Le Digabel, M. Forcato, S. Bicciato, N. Elvassore, S. Piccolo, Role of YAP/TAZ in mechanotransduction. Nature 474, 179–183 (2011).

20. M. Hanna, H. Liu, J. Amir, Y. Sun, S. W. Morris, M. A. Siddiqui, L. F. Lau, B. Chaqour, Mechanical regulation of the proangiogenic factor CCN1/CYR61 gene requires the combined activities of MRTF-A and CREB-binding protein histone acetyltransferase. J Biol Chem 284, 23125–23136 (2009).

21. H. K. Vanyai, F. Prin, O. Guillermin, B. Marzook, S. Boeing, A. Howson, R. E. Saunders, T. Snoeks, M. Howell, T. J. Mohun, B. Thompson, Control of skeletal morphogenesis by the Hippo-YAP/TAZ pathway. Development 147, (2020).

22. H. Zhang, H. A. Pasolli, E. Fuchs, Yes-associated protein (YAP) transcriptional coactivator functions in balancing growth and differentiation in skin. Proc Natl Acad Sci U S A 108, 2270–2275 (2011).

23. N. Chen, C. C. Chen, L. F. Lau, Adhesion of human skin fibroblasts to Cyr61 is mediated through integrin alpha 6beta 1 and cell surface heparan sulfate proteoglycans. J Biol Chem 275, 24953–24961 (2000).

24. N. Chen, S. J. Leu, V. Todorovic, S. C. Lam, L. F. Lau, Identification of a novel integrin alphavbeta3 binding site in CCN1 (CYR61) critical for pro-angiogenic activities in vascular endothelial cells. J Biol Chem 279, 44166–44176 (2004).

25. S. J. Leu, Y. Liu, N. Chen, C. C. Chen, S. C. Lam, L. F. Lau, Identification of a novel integrin alpha 6 beta 1 binding site in the angiogenic inducer CCN1 (CYR61). J Biol Chem 278, 33801–33808 (2003).

26. J. M. Schober, L. F. Lau, T. P. Ugarova, S. C. Lam, Identification of a novel integrin alphaMbeta2 binding site in CCN1 (CYR61), a matricellular protein expressed in healing wounds and atherosclerotic lesions. J Biol Chem 278, 25808–25815 (2003).

27. A. M. Babic, M. L. Kireeva, T. V. Kolesnikova, L. F. Lau, CYR61, a product of a growth factor-inducible immediate early gene, promotes angiogenesis and tumor growth. Proc Natl Acad Sci U S A 95, 6355–6360 (1998).

28. L. F. Lau, CCN1/CYR61: the very model of a modern matricellular protein. Cell Mol Life Sci 68, 3149–3163 (2011).

29. T. M. Grzeszkiewicz, V. Lindner, N. Chen, S. C. Lam, L. F. Lau, The angiogenic factor cysteine-rich 61 (CYR61, CCN1) supports vascular smooth muscle cell adhesion and stimulates chemotaxis through integrin alpha(6)beta(1) and cell surface heparan sulfate proteoglycans. Endocrinology 143, 1441–1450 (2002).

30. T. P. O’Brien, L. F. Lau, Expression of the growth factor-inducible immediate early gene cyr61 correlates with chondrogenesis during mouse embryonic development. Cell Growth Differ 3, 645–654 (1992).

31. Y. Zhang, T. J. Sheu, D. Hoak, J. Shen, M. J. Hilton, M. J. Zuscik, J. H. Jonason, R. J. O’Keefe, CCN1 Regulates Chondrocyte Maturation and Cartilage Development. J Bone Miner Res 31, 549–559 (2016).

32. G. Zhao, B. L. Huang, D. Rigueur, W. Wang, C. Bhoot, K. R. Charles, J. Baek, S. Mohan, J. Jiang, K. M. Lyons, CYR61/CCN1 Regulates Sclerostin Levels and Bone Maintenance. J Bone Miner Res 33, 1076–1089 (2018).

33. G. Zhao, E. W. Kim, J. Jiang, C. Bhoot, K. R. Charles, J. Baek, S. Mohan, J. S. Adams, S. Tetradis, K. M. Lyons, CCN1/Cyr61 Is Required in Osteoblasts for Responsiveness to the Anabolic Activity of PTH. J Bone Miner Res 35, 2289–2300 (2020).

34. M. Hadjiargyrou, W. Ahrens, C. T. Rubin, Temporal expression of the chondrogenic and angiogenic growth factor CYR61 during fracture repair. J Bone Miner Res 15, 1014–1023 (2000).

35. J. Lienau, H. Schell, D. R. Epari, N. Schütze, F. Jakob, G. N. Duda, H. J. Bail, CYR61 (CCN1) protein expression during fracture healing in an ovine tibial model and its relation to the mechanical fixation stability. J Orthop Res 24, 254–262 (2006).

36. S. Ali, S. R. Hussain, A. Singh, V. Kumar, S. Walliullah, N. Rizvi, M. Yadav, M. K. Ahmad, A. A. Mahdi, Study of Cysteine-Rich Protein 61 Genetic Polymorphism in Predisposition to Fracture Nonunion: A Case Control. Genet Res Int 2015, 754872 (2015).

37. S. P. Frey, S. Doht, L. Eden, S. Dannigkeit, N. Schuetze, R. H. Meffert, H. Jansen, Cysteine-rich matricellular protein improves callus regenerate in a rabbit trauma model. Int Orthop 36, 2387–2393 (2012).

38. V. Röntgen, R. Blakytny, R. Matthys, M. Landauer, T. Wehner, M. Göckelmann, P. Jermendy, M. Amling, T. Schinke, L. Claes, A. Ignatius, Fracture healing in mice under controlled rigid and flexible conditions using an adjustable external fixator. Journal of Orthopaedic Research 28, 1456–1462 (2010).

39. J. D. Boerckel, B. A. Uhrig, N. J. Willett, N. Huebsch, R. E. Guldberg, Mechanical regulation of vascular growth and tissue regeneration in vivo. Proc Natl Acad Sci U S A 108, E674–680 (2011).

40. H.-J. Choi, H. Zhang, H. Park, K.-S. Choi, H.-W. Lee, V. Agrawal, Y.-M. Kim, Y.-G. Kwon, Yes-associated protein regulates endothelial cell contact-mediated expression of angiopoietin-2. Nature Communications 6, 6943 (2015).

41. J. Kim, Y. H. Kim, J. Kim, D. Y. Park, H. Bae, D. H. Lee, K. H. Kim, S. P. Hong, S. P. Jang, Y. Kubota, Y. G. Kwon, D. S. Lim, G. Y. Koh, YAP/TAZ regulates sprouting angiogenesis and vascular barrier maturation. J Clin Invest 127, 3441–3461 (2017).

42. X. Wang, A. Freire Valls, G. Schermann, Y. Shen, I. M. Moya, L. Castro, S. Urban, G. M. Solecki, F. Winkler, L. Riedemann, R. K. Jain, M. Mazzone, T. Schmidt, T. Fischer, G. Halder, C. Ruiz de Almodóvar, YAP/TAZ Orchestrate VEGF Signaling during Developmental Angiogenesis. Developmental Cell 42, 462–478.e467 (2017).

43. Q. Cong, Y. Liu, T. Zhou, Y. Zhou, R. Xu, C. Cheng, H. S. Chung, M. Yan, H. Zhou, Z. Liao, B. Gao, G. A. Bocobo, T. A. Covington, H. J. Song, P. Su, P. B. Yu, Y. Yang, A self-amplifying loop of YAP and SHH drives formation and expansion of heterotopic ossification. Science Translational Medicine 13, eabb2233 (2021).

44. C. A. Fullenkamp, S. L. Hall, O. I. Jaber, B. L. Pakalniskis, E. C. Savage, J. M. Savage, G. K. Ofori-Amanfo, A. M. Lambertz, S. D. Ivins, C. S. Stipp, B. J. Miller, M. M. Milhem, M. R. Tanas, TAZ and YAP are frequently activated oncoproteins in sarcomas. Oncotarget 7, 30094–30108 (2016).

45. M. R. Tanas, A. Sboner, A. M. Oliveira, M. R. Erickson-Johnson, J. Hespelt, P. J. Hanwright, J. Flanagan, Y. Luo, K. Fenwick, R. Natrajan, C. Mitsopoulos, M. Zvelebil, B. L. Hoch, S. W. Weiss, M. Debiec-Rychter, R. Sciot, R. B. West, A. J. Lazar, A. Ashworth, J. S. Reis-Filho, C. J. Lord, M. B. Gerstein, M. A. Rubin, B. P. Rubin, Identification of a disease-defining gene fusion in epithelioid hemangioendothelioma. Sci Transl Med 3, 98ra82 (2011).

46. M. L. Kireeva, F. E. Mo, G. P. Yang, L. F. Lau, Cyr61, a product of a growth factor-inducible immediateearly gene, promotes cell proliferation, migration, and adhesion. Mol Cell Biol 16, 1326–1334 (1996).

47. A. Hasan, N. Pokeza, L. Shaw, H. S. Lee, D. Lazzaro, H. Chintala, D. Rosenbaum, M. B. Grant, B. Chaqour, The matricellular protein cysteine-rich protein 61 (CCN1/Cyr61) enhances physiological adaptation of retinal vessels and reduces pathological neovascularization associated with ischemic retinopathy. J Biol Chem 286, 9542–9554 (2011).

48. C. C. Chen, F. E. Mo, L. F. Lau, The angiogenic factor Cyr61 activates a genetic program for wound healing in human skin fibroblasts. J Biol Chem 276, 47329–47337 (2001).

49. D. Zhou, D. J. Herrick, J. Rosenbloom, B. Chaqour, Cyr61 mediates the expression of VEGF, alphavintegrin, and alpha-actin genes through cytoskeletally based mechanotransduction mechanisms in bladder smooth muscle cells. J Appl Physiol (1985) 98, 2344–2354 (2005).

50. H. Duan, Z. He, M. Lin, Y. Wang, F. Yang, R. A. Mitteer, H. J. Kim, E. Yeo, H. Han, L. Qin, Y. Fan, Y. Gong, Plasminogen regulates mesenchymal stem cell-mediated tissue repair after ischemia through Cyr61 activation. JCI Insight 5, (2020).

51. L. Wang, L. Yao, H. Duan, F. Yang, M. Lin, R. Zhang, Z. He, J. Ahn, Y. Fan, L. Qin, Y. Gong, Plasminogen Regulates Fracture Repair by Promoting the Functions of Periosteal Mesenchymal Progenitors. J Bone Miner Res 36, 2229–2242 (2021).

52. M. Wong, M. L. Kireeva, T. V. Kolesnikova, L. F. Lau, Cyr61, product of a growth factor-inducible immediate-early gene, regulates chondrogenesis in mouse limb bud mesenchymal cells. Dev Biol 192, 492–508 (1997).

53. H. Liu, F. Peng, Z. Liu, F. Jiang, L. Li, S. Gao, G. Wang, J. Song, E. Ruan, Z. Shao, R. Fu, CYR61/CCN1 stimulates proliferation and differentiation of osteoblasts in vitro and contributes to bone remodeling in vivo in myeloma bone disease. Int J Oncol 50, 631–639 (2017).

54. W. Si, Q. Kang, H. H. Luu, J. K. Park, Q. Luo, W. X. Song, W. Jiang, X. Luo, X. Li, H. Yin, A. G. Montag, R. C. Haydon, T. C. He, CCN1/Cyr61 is regulated by the canonical Wnt signal and plays an important role in Wnt3A-induced osteoblast differentiation of mesenchymal stem cells. Mol Cell Biol 26, 2955–2964 (2006).

55. J.-L. Su, J. Chiou, C.-H. Tang, M. Zhao, C.-H. Tsai, P.-S. Chen, Y.-W. Chang, M.-H. Chien, C.-Y. Peng, M. Hsiao, M.-L. Kuo, M.-L. Yen, CYR61 Regulates BMP-2-dependent Osteoblast Differentiation through the αvβ3 Integrin/Integrin-linked Kinase/ERK Pathway*. Journal of Biological Chemistry 285, 31325–31336 (2010).

56. I. Chiesa, C. De Maria, A. Lapomarda, G. M. Fortunato, F. Montemurro, R. Di Gesù, R. S. Tuan, G. Vozzi, R. Gottardi, Endothelial cells support osteogenesis in an in vitro vascularized bone model developed by 3D bioprinting. Biofabrication 12, 025013 (2020).

57. A. Anup, S. Dieterich, R. O. C. Oreffo, H. L. Dailey, A. Lang, M. Haffner-Luntzer, K. R. Hixon, Embracing ethical research: Implementing the 3R principles into fracture healing research for sustainable scientific progress. Journal of Orthopaedic Research n/a, (2023).

58. P. Jirkof, M. Durst, R. Klopfleisch, R. Palme, C. Thöne-Reineke, F. Buttgereit, K. Schmidt-Bleek, A. Lang, Administration of Tramadol or Buprenorphine via the drinking water for post-operative analgesia in a mouse-osteotomy model. Scientific Reports 9, 10749 (2019).

59. A. Lang, A. Schulz, A. Ellinghaus, K. Schmidt-Bleek, Osteotomy models - the current status on pain scoring and management in small rodents. Lab Anim 50, 433–441 (2016).

